# The association between pain-induced autonomic reactivity and descending pain control is mediated by the periaqueductal grey

**DOI:** 10.1101/2020.10.26.355529

**Authors:** Elena Makovac, Alessandra Venezia, David Hohenschurz-Schmidt, Ottavia Dipasquale, Jade B Jackson, Sonia Medina, Owen O’Daly, Steve CR Williams, Stephen B McMahon, Matthew A Howard

## Abstract

There is a strict interaction between the autonomic nervous system (ANS) and pain, which might involve descending pain modulatory mechanisms. The periaqueductal grey (PAG) is involved both in descending pain modulation and ANS, but its role in mediating this relationship has not yet been explored.

Here, we sought to determine brain regions mediating ANS and descending pain control associations. 30 participants underwent Conditioned Pain Modulation (CPM) assessments, in which they rated painful pressure stimuli applied to their thumbnail, either alone or with a painful cold contralateral stimulation. Differences in pain ratings between ‘pressure-only’ and ‘pressure+cold’ stimuli provided a measure of descending pain control. In 18 of the 30 participants, structural scans and two functional MRI assessments, one pain-free and one during cold-pain, were acquired. Heart Rate Variability (HRV) was simultaneously recorded.

Low frequency HRV (LF-HRV) and the CPM score were negatively correlated; individuals with higher LF-HRV during pain reported reductions in pain during CPM. PAG-frontal medial cortex (FMC) and PAG-rostral ventro-medial medulla (RVM) functional connectivity correlated negatively with the CPM. Importantly, PAG-FMC functional connectivity mediated the strength of HRV-CPM association. CPM response magnitude was also negatively associated with PAG and positively associated with FMC grey matter volumes.

Our multi-modal approach, using behavioral, physiological and MRI measures, provides important new evidence of interactions between ANS and descending pain mechanisms. ANS dysregulation and dysfunctional descending pain modulation are characteristics of chronic pain. We suggest that further investigation of body-brain interactions in chronic pain patients may catalyse the development of new treatments.

## Introduction

Nociception is modulated by descending pain control systems in the brainstem and spinal cord, under the direct influence of higher order brain areas. In humans, the efficiency of these systems can be measured using paradigms such as Conditioned Pain Modulation (CPM) (Yarnitsky, 2010). During a typical CPM trial, participants subjectively rate responses to a painful test stimulus, both on its own, and in the presence of an additional painful conditioning stimulus (Yarnitsky et al., 2015). Here, a reduction in the pain induced by the test stimulus when coincident with the conditioning stimulus indicates efficient descending pain control. CPM responses are, however, variable, and a reduction in pain during conditioning stimulus application is only observed in some individuals (Kemp, Kennedy, Wu, Ridout, & Rice, 2019). Individual differences in CPM responses may relate to factors such as age or gender (Edwards, Ness, Weigent, & Fillingim, 2003; Granot et al., 2008), emotions (Roy, Piché, Chen, Peretz, & Rainville, 2009), personality (Nahman-Averbuch, Yarnitsky, Sprecher, Granovsky, & Granot, 2016) or the type of paradigm adopted (Kemp et al., 2019). Despite substantial interindividual variability, reports indicate CPM responses predict post-operative pain (Yarnitsky et al., 2008) and pharmacological treatment response (Yarnitsky, Granot, Nahman-Averbuch, Khamaisi, & Granovsky, 2012), thus showing potential as a biomarker.

The bi-directional interaction between the cardiovascular system and pain is another important component of descending pain control (Bruehl & Chung, 2004). In healthy individuals, acute pain increases sympathetic arousal and blood pressure (Kyle & McNeil, 2014). By contrast, a reduction in pain perception is reported in healthy participants during spontaneous or induced high blood pressure (Saccò et al., 2013). Similarly, unmedicated patients with hypertension report lower pain sensitivity and higher pain thresholds, a condition termed as blood pressure-related hypoalgesia (Olsen et al., 2013; Ottaviani et al., 2018). Several studies have suggested that high blood pressure may protect against clinical pain (Bruehl, Chung, Ward, Johnson, & McCubbin, 2002; France & Katz, 1999; Hagen et al., 2005; Maixner, Fillingim, Kincaid, Sigurdsson, & Harris, 1997), entertaining the possibility that the baroreflex -a homeostatic mechanism that controls blood pressure-may also be involved in pain regulation (Suarez-Roca et al., 2019). An association has been also described between blood pressure and the CPM response (Chalaye, Lafrenaye, Goffaux, & Marchand, 2014), supporting the role of baroreceptors that signal the cardiovascular state of the body to the brain in pain control (Suarez-Roca et al., 2018). Carotid baroreceptor stimulation results in reduced pain sensitivity in both hypertensive and normotensive individuals (L. Edwards et al., 2003), suggesting that interactions between the ANS and descending pain control systems may involve baroreflex activation (Dayan et al., 2018). Heart rate variability (HRV), a measure of an individual sympatho/vagal balance, has also been shown to be related to the perception of pain. Increased low-frequency HRV, a measure specifically linked to baroreflex efficiency (Moak et al., 2007; Rahman, Pechnik, Gross, Sewell, & Goldstein, 2011) has been associated with reduced subjective ratings of unpleasantness to cold pain and higher thresholds for low-to moderate levels of thermal pain (Appelhans & Luecken, 2008).

Perhaps associations between the ANS and pain control are not surprising, given the overlap between brain structures involved in ANS regulation (via the baroreflex) and those involved in pain. For example, the frontal medial (FMC) and prefrontal cortex (PFC), anterior cingulate cortex, thalamus and insula are involved both in the regulation of body autonomic arousal (Beissner, Meissner, Bär, & Napadow, 2013; Benarroch, 2012; Sklerov, Dayan, & Browner, 2018) and in pain perception and elaboration (Benarroch, 2006; Peyron, Laurent, & Garcia-Larrea, 2000). Animal studies indicate that the periaqueductal grey (PAG) and the rostral ventromedial medulla (RVM) have well-defined roles in descending pain control (Bouhassira, Villanueva, Bing, & le Bars, 1992; Harris, 1996; Villanueva, Bouhassira, & Le Bars, 1996). In humans, pain control mechanisms have been related to functional networks involving the PAG and the FMC/PFC (Bogdanov et al., 2015) and to resting-state functional connectivity (FC) between the PAG and RVM (Harper et al., 2018; Yelle, Oshiro, Kraft, & Coghill, 2009); but also between the subnucleus reticularis dorsalis and cingulate and dorsolateral prefrontal areas (Youssef, Macefield, & Henderson, 2016). These same brain structures namely, the cingulate cortex, insula, amygdala, and PAG are also associated with reduced baroreflex sensitivity, particularly the ability of the baroreflex to control transient BP fluctuations in response to physiological stress (Gianaros, Onyewuenyi, Sheu, Christie, & Critchley, 2012).

The neurovisceral integration model (NVI) (Thayer and Lane, 2000, 2007) provides a framework for investigating inter-relationships between descending pain control mechanisms and autonomic functioning in the brain. The NVI model theorises that increased HRV provides an individual with flexibility to adapt in response to changing physiological and environmental demands, governed by a ‘Central Autonomic Network’ (CAN) (Benarroch, 1993), incorporating the prefrontal cortex (anterior cingulate, insula, orbitofrontal, and ventromedial cortex), amygdala, hypothalamus, and brain stem (including the PAG and RVM). The dynamic and bi-directional interactions between the CAN and the cardiovascular system affect cognition, executive functioning and emotion (Thayer, Hansen, Saus-Rose, & Johnsen, 2009).

Here, we sought to determine brain mechanisms underlying the putative association between descending pain control and autonomic regulation systems in a group of healthy participants. We employed resting-state fMRI (rs-fMRI), and voxel-based morphometry (VBM) as measures of brain function and structure, respectively, focusing our attention on the PAG as a major hub for both autonomic and pain processing. We also examined HRV as an index of autonomic reactivity to pain and performed a behavioral experiment examining descending pain control. We hypothesised that connectivity between the PAG and other pain-related structures would mediate the association between the ANS and descending pain control mechanisms. We hypothesised an involvement of the RVM and FMC, given their role in descending pain controls, as documented by animal studies (REF) recent studies in healthy volunteers and chronic pain patients (Harper et al., 2018; Moont, Crispel, Lev, Pud, & Yarnitsky, 2011).

## METHODS AND MATERIALS

### Participants

30 healthy participants took part in the CPM experiment (12 women, 18 men, mean age across the group = 26.4; SD = 4.7 years). Out of these 30 participants, only 18 (6 women, 12 men, mean age across the group = 27.1; SD = 5.1 years) took part in both the CPM and the MRI and physiology experimental session (detailed below). Therefore, 30 participants in total were considered for the behavioural (CPM) analyses whilst 18 participants only were considered for the regression analysis with MRI and HRV data. All participants were righthanded as assessed by the Edinburgh handedness inventory (Oldfield, 1971). Exclusion criteria included MRI contraindications, history of brain injuries, hypertension, neurological or psychiatric disease, and alcohol or drug abuse. To minimise the potential effects of menstrual cycle-related hormone fluctuations on pain responses (Vincent & Tracey, 2010) and descending pain modulating effects (Tousignant-Laflamme & Marchand, 2009), female participants were all tested within the follicular phase of their menstrual cycle. Furthermore, to minimise the influence of diurnal variations on pain responses (Strian, Lautenbacher, Galfe, & Hölzl, 2016) and on rs-fMRI networks activity (Jiang et al., 2016), participants were always tested at the same time during the day across the three experimental sessions. At the beginning of each visit, participants were tested for drug use (urine drug test) and alcohol consumption (alcohol breathalyser). All participants provided written informed consent. The study was approved by the King’s College London Research Ethics Committee (HR-16/17-4769).

### Experimental procedure

Participants took part in one familiarisation session, one scanning session and one post-scanning CPM session. At the beginning of each experimental session, participants completed the ‘state’ subscale of the State-Trait Anxiety Inventory (STAI) (Spielberger, Gorsuch, Lushene, Vagg, & Jacobs, 1983), in order to evaluate for possible differences in anxiety levels between the three testing sessions.

### Familiarisation and thresholding session

During the familiarisation session, participants became accustomed to the neuroimaging environment (in a simulated ‘mock’ scanner), and with the cold-pain stimulation paradigm. The cold pain stimulation was delivered via an aluminium probe (4 x 20 cm), attached to the volar surface of their right forearm, through which cold water (2°C) was constantly circulated by means of two chillers.

Next, participants were individually thresholded for pressure pain. A laboratory-built pressure-inducing thumb device was used to apply pressure on the right-side thumbnail (see Supplementary material Figure S1 for a picture of the experimental device). Participants received an ascending series of pressure stimuli and were asked to state when they felt the first sensation of pain (pain threshold) and when they reached a 70 on a verbal scale from 0 (no pain) to 100 (maximum pain tolerance). Three intermediate pressure values between these thresholds were created to provide five equally spaced pressure values, starting with the threshold and ending with the highest values corresponding to each individuals’ VAS 70/100. For example, if the minimum pain threshold was represented by a pressure of 200 kPa in the ascending series and the maximum pain threshold of > 70 was reached with a pressure of 600 kPa, the randomised series would consist of pressures of 200, 300, 400, 500 and 600 kPa. The five pressure values were then presented to the participant in a random order, each repeated three-times (total number of new stimulations n = 15). After each stimulation, the participant was required to give a rating on a VAS scale. Based on the 15 pressure values and 15 subjective pain ratings, a polynomial regression function was used to determine each individual’s representation of VAS 50, 60 and 70 (Kosek et al., 2017). These values were then used in our CPM paradigm (third session).

#### MRI scanning session and physiological recording

During the scanning sessions, participants underwent three FC MRI scans, each of six minutes in duration: a baseline; a cold-pain and a post-cold recovery block. The investigation of the post-cold recovery phase aimed at identifying pain-related carry over effects and the results from this analysis have been reported elsewhere (Makovac et al., 2019). A multi-echo resting-state fMRI (rs-fMRI) approach was adopted to mitigate problems related to pain-induced movements, which are an important source of non-BOLD artifacts in pain studies (Dipasquale et al., 2017; Moayedi, Salomons, & Atlas, 2018; Siegel et al., 2016). During each period, participants were instructed to keep still with their eyes open, focusing on a fixation cross presented at the center of the screen, without thinking of anything or falling asleep.

MRI images were acquired on a GE MR750 scanner, equipped with a 32-channel receive-only head coil (NovaMedical). Structural volumes were obtained using a high-resolution three-dimensional magnetization-prepared rapid gradient-echo sequence (TR = 7312 ms, TE = 3.02 ms, flip angle = 11°, slice thickness = 12 mm, 196 sagittal slices, FOV = 270 mm). Multi-echo rs-fMRI datasets were acquired using a T2*-weighted blood oxygenation level dependent (BOLD) sensitive echo planar imaging (EPI) sequence (TR = 2 s, TE1 = 12 ms, TE2 = 28 ms; TE3= 44 ms; flip-angle 80°, 32 slices, 3mm slice thickness, 240 mm FOV, voxel size 3.75 x 3.75 x 3 mm). During rs-fMRI, heart rate was monitored using an inbuilt MRI-compatible finger pulse oximeter (General Electric) recorded digitally from an analogue output waveform at a sample rate of 40 Hz, fitted to the participants’ dominant right index finger.

#### CPM session

Participants were tested during a third session (on a different day from the scanning session), where they underwent a CPM paradigm, in order to test descending pain modulatory mechanisms.

##### CPM experimental paradigm

During CPM, short-duration noxious pressure test stimuli were delivered to the righthand thumbnail prior to, simultaneously with, and immediately following a noxious cold conditioning stimulus applied to the left-hand forearm. The intensity of the pressure test stimuli was determined individually for each participant, based on the previously described thresholding procedure, to synthesise pressures to induce pain ratings of 50, 60 and 70 on a 0-100 VAS. Given the lack on unanimous consensus regarding the optimal psychophysical characteristics of the test stimulus and conditioning stimulus in eliciting the CPM effect, we randomly delivered pressure pain stimuli corresponding to the three subjective intensities (50, 60 and 70 VAS), to explore the effect of test stimulus intensity on the CPM response. To account for potential carry-over effects of the conditioning stimulus, test stimuli were presented simultaneously with the conditioning stimulus, and for 6 minutes following the conditioning stimulus.

Participants underwent three separate experimental runs (refer to Figure 1); i) test stimuli only (Baseline condition), and two identical CPM runs (CPM1 and CPM2), separated by a fifteen-minute break. CPM investigations were split into two runs to provide an adequate number of events per experimental condition but to minimise response habituation and/or peripheral sensitisation. The CPM runs comprised two six-minute mini-blocks, the first during which cold stimulation was present (CPM Cold-ON condition), the second without cold stimulation (CPM Cold-OFF condition). During the Baseline condition, participants rated the intensity of pain evoked by pressure pain stimuli, on a computerized VAS ranging from 0 (no pain) to 100 (maximal imaginable pain). Twenty-four pressure test stimuli were delivered in a random order to their right thumbnail, eight per class of stimulus intensity (50, 60 and 70 VAS). Each test stimulus had a duration of two seconds, followed by an eight-second VAS rating period then five-second inter-trial interval. The total duration of the Baseline run was 9.2 minutes. During the CPM run (CPM1 and CPM2), participants were instructed to rate the pressure pain test stimuli (as per the Baseline condition), and not the painful cold conditioning stimulus applied to their left forearm. Finally, participants subjectively rated the intensity of the painful cold conditioning stimulus using a computerized VAS ranging from 0 (no pain) to 100 (maximal imaginable pain).

**Figure 1.**
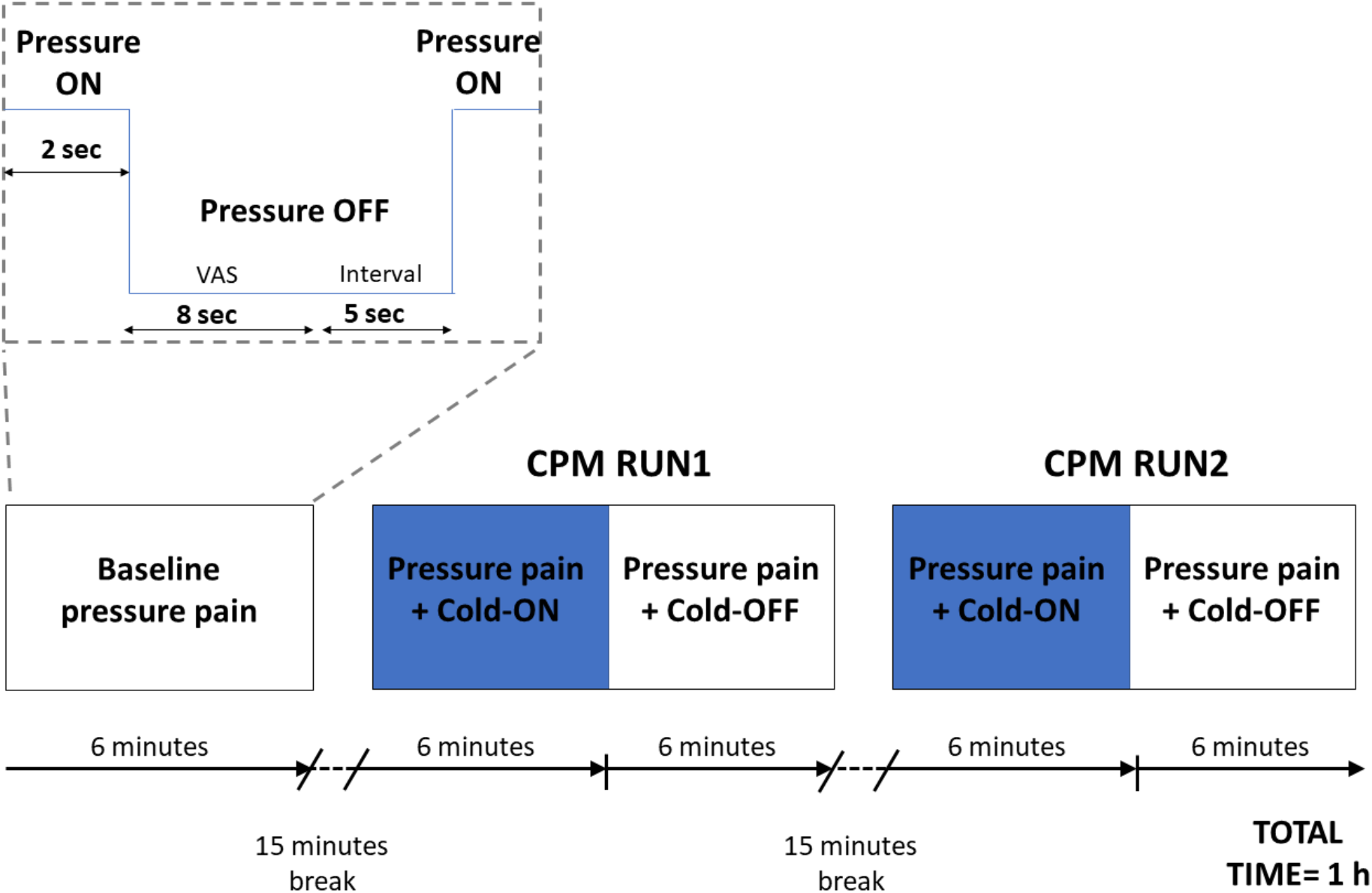
A graphical representation of the experimental paradigm. Participants were required to rate 2-seconds painful pressure stimuli for 6 minutes (Baseline condition), followed by a 15-minute break. During CPM, participants rated pressure pain with a simultaneous contralateral cold stimulation for 6 minutes (CPM Cold-ON condition), followed by ratings of pressure pain without the contralateral cold (CPM Cold-OFF condition).

### Statistical analysis

#### State anxiety

A one-way within-subject ANOVA was used to explore for differences in state anxiety among the three experimental sessions (thresholding, scanning session, CPM session). Possible associations between state anxiety, HRV and CPM were investigated by means of Pearson’s correlation coefficient. Statistical threshold was set at p < 0.05.

#### CPM analysis

CPM Cold-ON and CPM Cold-OFF conditions were obtained by averaging pressure ratings from CPM run 1 and CPM run 2 (Figure 1). ΔCPM Cold-ON and ΔCPM Cold-OFF scores were calculated by subtracting Baseline VAS rating values from the CPM Cold-ON and CPM Cold-OFF conditions ratings, respectively. Negative ΔCPM values indicated a decrease in perceived pressure pain intensity during the conditioning stimulus (ΔCPM Cold-ON) or following the conditioning stimulus (ΔCPM Cold-OFF), whereas positive values indicated an increase in perceived pain. Participants in whom ΔCPM Cold-ON and ΔCPM Cold-OFF scores were 2 standard errors of the mean (s.e.m.) above or below the group average were classified as facilitators (i.e. the conditioning stimulus resulted in an increase in the perceived pressure pain) or inhibitors (i.e. the conditioning stimulus resulted in an inhibition of the perceived pressure pain), respectively. ΔCPM values were used for further correlational and fMRI analyses.

A two-way within-subject ANOVA was used to explore the main effect of Pressure (50,60 and 70 VAS), Condition (Baseline, CPM Cold-ON, CPM Cold-OFF), and the Pressure x Condition interaction.

#### Heart Rate Variability analysis

Inter-beat interval values were visually inspected, and potential artefacts were manually removed. Inter-beat-intervals were then entered into Kubios HRV Standard ver. 3.0.2 (Tarvainen et al., 2014) and further artefact correction was applied using the lowest correction threshold appropriate for individual artefact severity. In stationary conditions, pulse oximetry has been shown to be a good surrogate measure for ECG-derived HRV (Gil et al., 2010; Schäfer & Vagedes, 2013). To capture the most salient autonomic response to pain (without the interfering effect of habituation), low-frequency HRV (LF-HRV; 0.04–0.15 Hz) was derived from recordings of the initial 3 minutes during cold-pain application. LF-HRV is a measure of particular interest in acute pain studies as it is an index of baroreflex activity (Goldstein, Bentho, Park, & Sharabi, 2011; Rahman et al., 2011). Recent meta-analytic works have highlighted the influence of acute experimental pain on LF-HRV (Koenig, Jarczok, Ellis, Hillecke, & Thayer, 2014).

Next, LF-HRV data were normalised. Normalised units (nu) of HRV spectral data express the relative value of each frequency component in relation to the total power spectrum minus the very low frequency band (VLF): *LF (nu)* = *LF (ms^2^) / (total power (ms^2^) - VLF (ms^2^)* (Tarvainen et al., 2014).

#### Rs-fMRI preprocessing

AFNI (Cox, 1996), the Advanced Normalization Tools (ANTs) (Avants et al., 2011) and FSL (Smith et al., 2004) were used to process multi-echo rs-fMRI data. The volumes were first re-aligned and slice timing-corrected. Next, multi-echo sequences were optimally combined (OC) by taking a weighted summation of the three echoes using an exponential T2* weighting approach (Posse et al., 1999). A multi-Echo ICA approach (implemented by the tool meica.py, Version v2.5 beta9) (P; Kundu et al., 2013; P. Kundu et al., 2013) was adopted to de-noise the OC-data. Multi-Echo ICA is an effective method for the removal of physiological and motion-related noise, resulting in a significant increase in the temporal SNR (Dipasquale et al., 2017; P. Kundu et al., 2013). Briefly, multi-echo principal component analysis was first used to reduce the data dimensionality in the OC dataset. Spatial ICA was then applied on one echo, and the independent component time-series were fitted to the pre-processed time-series from each of the three echoes to generate ICA weights for each echo. These weights were then fitted to the linear TE-dependence and TE-independence models to generate F-statistics and component-level κ and ρ values, which respectively indicate BOLD and non-BOLD weightings. The κ and ρ metrics were then used to identify non-BOLD-like components to be regressed out of the OC dataset as noise. For further technical details on ME-ICA see (Kundu et al., 2015). Lastly, the white matter (WM) and cerebrospinal fluid (CSF) signals were regressed-using FSL. A high-pass temporal filter (with a cut-off frequency of 0.005 Hz) and spatial smoothing with a 5 mm FWHM Gaussian kernel were then applied. Each participant’s dataset was co-registered to its corresponding structural scan with an affine registration and normalised to standard MNI152 space (with a non-linear approach) with a 2×2×2mm^3^ resampling using ANTs.

#### VBM pre-processing

T1 weighted volumes from all participants were visually reviewed to exclude the presence of macroscopic artefacts, prior to preprocessing in SPM12 (Statistical Parametrical Mapping, http://www.fil.ion.ucl.ac.uk/spm/) using the DARTEL algorithm. In brief, the MR images were bias-field corrected to correct non-uniformities then segmented into grey matter, white matter and cerebro-spinal fluid sections using a priori tissue probability maps in International Consortium of Brain Mapping (ICBM) template space. The individual grey matter images were subsequently normalized to the Montreal Neurological Institute (MNI) template with a 1.5×1.5×1.5 mm3 voxel size and modulated to maintain original individual grey matter volumes. Finally, all grey matter images were smoothed with 8-mm full-width at half-maximum isotropic Gaussian kernel. Total intra-cranial volume (ICV) was calculated and used as covariate of no interest in subsequent GLM analyses (see VBM Analyses).

#### Seed-Based fMRI: First Level Analysis

An ROI for the PAG (a 3 mm-radius sphere; MNI = 0 −32 −10; (Coulombe et al., 2017)) was constructed using the Marsbar toolbox implemented in SPM 12 (http://marsbar.sourceforge.net/). The average resting state fMRI time-series for the ROI was extracted from smoothed images for each participant and scan and used as a regressor in a 1^st^ level SPM analysis, to explore the network of areas associated with the seed region.

#### Assessment of Relationships Between Brain FC, Brain Structure and Self-Reported Pain and HRV measures

For both seed-based fMRI and VBM analyses, we sought to determine brain regions positively or negatively associated with ΔCPM Cold-ON, with LF-HRV in response to cold, or with the interaction between the two variables (i.e. the interaction term obtained by multiplying ΔCPM Cold-ON with LF-HRV). We further explored the association between brain structure/FC and both pressure pain threshold and cold pain ratings to determine whether our results were dependent on descending pain modulating pathway or rather on more general characteristics of the stimuli (i.e. pressure pain threshold or subjective perception of cold pain) adopted in our paradigm.

##### Seed-based Second Level Analyses

For the second-level seed-based regression analyses, we calculated the ΔFC of the PAG network in response to cold pain, by subtracting baseline FC maps from those during cold pain. ΔFC maps were then used for regression analyses within the framework of the general linear model with ΔCPM, LF-HRV and the ΔCPM x LF-HRV interaction term as covariates of interest; these analyses were performed to explore brain areas in which FC during pain was associated with descending pain modulation and with the interaction between descending pain modulation and autonomic reactivity to pain.

##### VBM Second Level Analyses

Statistical analyses were performed on smoothed grey matter maps within the framework of the general linear model. Similarly to the seed-based fMRI analysis, ΔCPM, LF-HRV (during cold) and the ΔCPM x LF-HRV interaction term were used as variables of interest in regression analyses, whereas intra-cranial-volume volume was used as a nuisance covariate to consider potential confounding effects of individual differences in head size.

###### Small Volume Correction Analyses For a priori-Hypothesized Regions of Interest

To test a-priori hypotheses, small volume correction (SVC) analyses were performed in the FMC, PAG (for the structural analyses only) and rostral ventromedial medulla (RVM). A 5mm-radius spherical mask was created for the PAG (MNI= 0 −30 −12). For the RVM, three separate 3-mm radius spheres were created, centred at MNI (x,y) = [0 −34], at 3 contiguous rostro-caudal levels from MNI (z) = [−53 to −49], as described in (Marciszewski et al., 2018). The three contiguous spheres were then merged resulting in a final single ROI.Finally, an anatomical mask was defined for the FMC, derived from the SPM-12 anatomy toolbox(Eickhoff et al., 2005).

For each participant, we extracted grey matter volume values and FC parameter estimates from the FMC, PAG and RVM masks. These values were used for further mediation analyses. Grey matter volume values and FC parameter estimates were also extracted from significant voxels and plotted for illustrative purposes (and to explore for possible outliers). Parameter estimates from significant voxels were also used for additional correlation analysis, to investigate whether ΔCPM related brain areas were associated with LF-HRV and vice-versa.

###### Statistical Inference

For both types of analyses, statistical thresholds were set to p < 0.05 - FWE-corrected at cluster level (Worsley & Friston, 1995) (cluster size defined using uncorrected voxel-level threshold p < 0.001). The results were also examined using a less stringent cluster forming threshold (p<0.005), to minimize the risk of type II error.

#### Correlation and Mediation analyses

Mediation analysis was performed to ascertain whether pain and ANS related brain regions mediated the relationship between autonomic reactivity to pain and CPM response. Data analysis was performed with SPSS 22.0 for Windows (SPSS Inc, USA). First, Pearson’s correlation coefficient was used to explore the association between LF-HRV during cold and ΔCPM ratings (ΔCPM = [pressure + cold] - [pressure only baseline]). Next, the mean parameter estimates were extracted from SVC masks (PAG, RVM and FMC) were entered in a mediation analysis using PROCESS (Hayes, 2012) in IBM SPSS version 23.0 (IBM Corp., Armonk, NY). LF-HRV was introduced as an independent variable, ΔCPM was the outcome variable, and brain functional and structural parameter estimates from SVC masks were possible mediators. The indirect effect was tested with 20,000 percentile-corrected bootstrap confidence intervals (95%).

## RESULTS

### Subjective pain ratings following cold pain

Following six-minute cold stimulation, participants gave an average pain rating of 44.94 (SD= 21.82) on a 0 – 100 VAS scale. At the end of the six-minute post-cold pain recovery phase, the participant pain intensity (rated on the VAS) was 1.50 (SD = 3.37).

### Effect of anxiety

We did not observe a significant difference in state anxiety between the three experimental sessions (F(2,48)= 1.03, p=0.37).

### VAS ratings during CPM

11 participants showed an inhibiting effect of the conditioning stimulus (i.e. ΔCPM was 2 s.e.m lower than the group average), 15 participants showed a facilitating effect of the conditioning stimulus (i.e. ΔCPM was 2 s.e.m higher than the group average), whereas four participants were non-responders (i.e. their ΔCPM was less than 2 s.e.m. from the group average, Figure 2).

**Figure 2.**
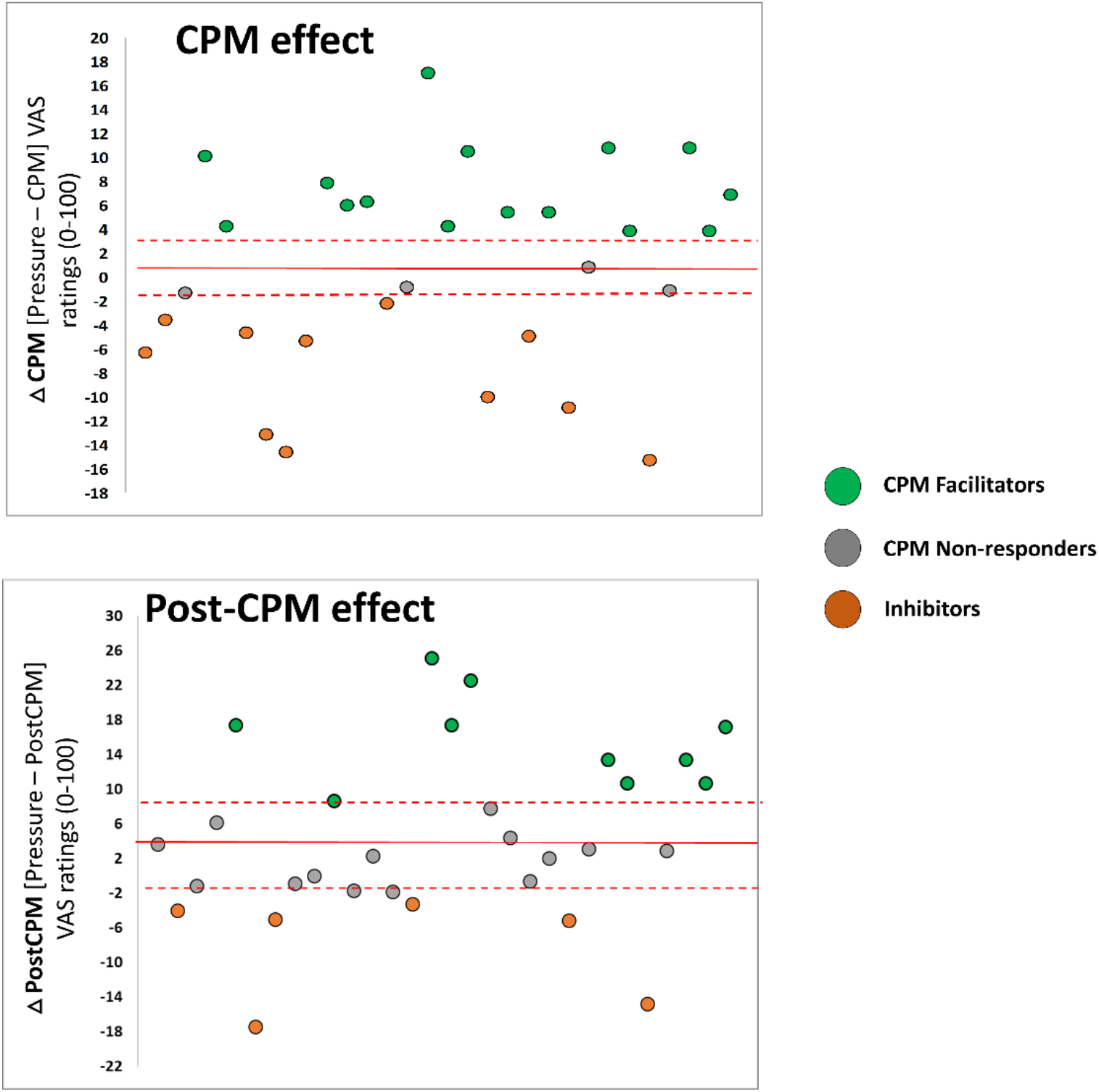
Conditioned pain modulation behavioural results. The magnitude of the CPM Cold-ON effect (Δ CPM Cold-ON) and CPM Cold-OFF effect (Δ CPM Cold-OFF) was defined as the difference in pressure pain rating between baseline (pressure only) and the CPM Cold-ON condition (cold conditioning stimulus simultaneously presented with pressure pain test stimulus) or CPM Cold-OFF condition (pressure pain test stimulus presented immediately following the cold conditioning stimulus). The red line represents the group average (+/- 2 s.e.m; dashed red lines). Overall, in the CPM Cold-ON condition, 11 participants showed an inhibiting effect (negative CPM values), 15 participants showed a facilitating effect of the conditioning stimulus (positive CPM values), whereas four participants were non-responders. During the CPM Cold-OFF condition, 6 participants presented an inhibiting effect, 10 participants showed a facilitating effect, whereas 14 participants were classified as non-responders.

A within-subject ANOVA analysis with two factors [Pressure (50, 60, 70 VAS)] and [ Condition (Baseline, Cold-ON, Cold-OFF)] was performed. A main effect of Pressure was found (F(2,58)= 4.77, p= 0.016), driven by higher VAS ratings in response to 60-pressure stimuli comparing to 50-pressure stimuli [mean (SD) = 31.5 (5.8) vs 38.3 (16.9) for 50 and 60 pressure respectively, t(20)= 6.92, p< 0.001]; higher ratings to 70-pressure stimuli [mean (SD)= 45.8 (18.2)] when compared to both 60-pressure [mean (SD)= 40.0 (18.2); t(29)= 5.87, p< 0.001] and 50-pressure stimulations [mean (SD)=34.3 (17.5); t(29)= 8.27, p< 0.001]. A main effect of Condition was also observed [F(2,58)= 34.31, p< 0.001], driven by an increase in overall perceived pain intensity in the CPM Cold-OFF condition [mean (SD)= 42.8 (19.8)] when compared to Baseline [mean (SD)=38.4 (17.3), t(29)= 2.43 p= 0.02] and CPM Cold-ON trials [mean (SD)=38.9 (17.9), t(29)= 3.0 p= 0.005]. No significant difference was observed when comparing CPM to Baseline [t<1].

During the CPM Cold-OFF condition, 6 participants showed an inhibiting effect, 10 participants showed a facilitating effect whereas 14 were classified as non-responders.

### Exploratory correlation analyses

Exploratory analyses were performed to investigate whether the magnitude of the CPM response was related to either the characteristics of the test stimulus, the conditioning stimulus, or to participants’ anxiety levels. Results indicated that the magnitude of the CPM Cold-ON response (ΔCPM Cold-ON) was not associated with cold pain ratings (p = 0.9), with individual pressure pain thresholds (p = 0.8), or state anxiety levels during the CPM session (p = 0.8).

### Correlation between HRV and CPM response

We observed a negative correlation between LF-HRV during cold pain and ΔCPM Cold-ON (r= −0.49, p=0.041), where individuals with the highest increase in LF-HRV during cold pain exhibited the strongest decrease in pressure pain rating during CPM (Figure 3). We did not observe a significant association between baseline LF-HRV and ΔCPM Cold-OFF.

**Figure 3.**
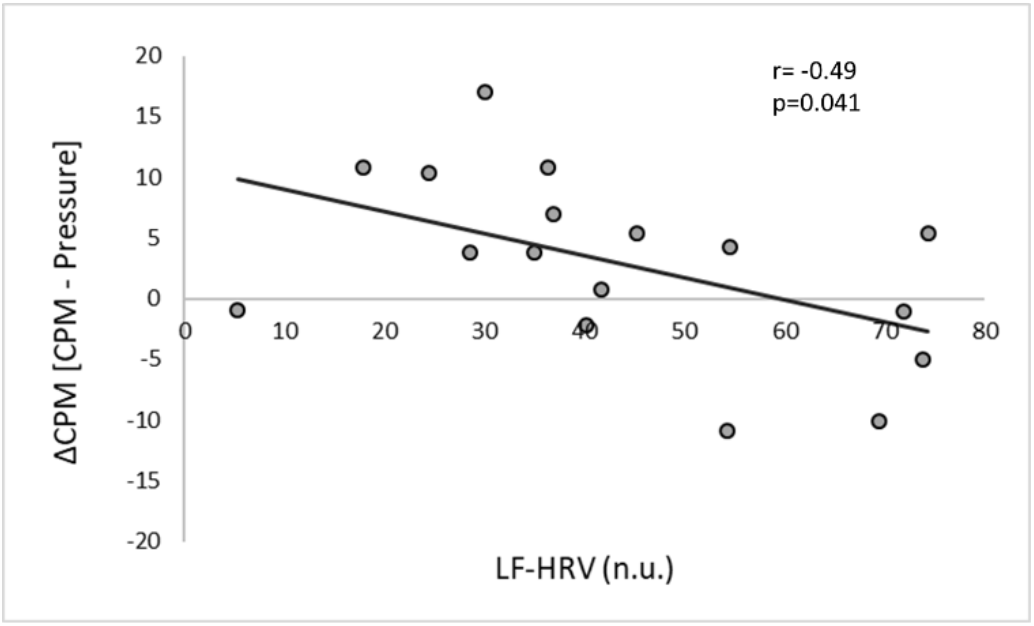
Negative correlation between ΔCPM Cold-ON [CPM Cold-ON–Baseline pressure pain]. and LF-HRV, indicating that individuals with the highest LF-HRV during cold pain exhibited a decrease in pressure pain rating during simultaneous contralateral cold pain. n.u. = normalized units.

### Seed-based analysis

Seed-based FC analysis of the PAG showed a network comprised of the bilateral parahippocampal/amygdalar areas, middle cingulate cortex, anterior insular cortex and prefrontal medial cortex in line with other published studies (see (Makovac et al., 2019).

### Associations between PAG functional connectivity, ΔCPM and autonomic reactivity (LF-HRV) during cold pain

#### Association between PAG FC and ΔCPM Cold-ON, LF-HRV and ΔCPM Cold-ON x LF-HRV interaction

A positive association was observed between ΔCPM scores and ΔFC in response to cold pain between the PAG and the supramarginal gyrus and left precuneus. At a more liberal cluster forming threshold (cluster forming threshold p<0.005) we also observed a negative association between ΔCPM Cold-ON and the PAG ΔFC in response to pain, with both the FMC and RVM (extending to the cerebellum). This result indicates that individuals with more efficient descending pain inhibition (as indexed by negative ΔCPM values) had a lower increase in FC between the PAG and more posterior parietal areas and stronger increase in FC between the PAG and the FMC and RVM during cold pain. At a more liberal cluster forming threshold (p<0.005), we observed a negative association between LF-HRV and the ΔFC between PAG and right angular gyrus (Table 2). We did not observe any association between PAG FC and the ΔCPM Cold-ON x LF-HRV interaction term.

#### Exploratory analyses of the association between PAG FC and pressure pain threshold and cold-pain ratings

We did not observe any positive or negative association between PAG ΔFC and baseline pressure pain ratings.

### Associations between brain structure, CPM score and autonomic reactivity (LF-HRV) during cold pain

We observed a negative correlation between ΔCPM Cold-ON score and FMC volume, and a positive correlation between ΔCPM Cold-ON and PAG volume (Figure 4, 2). These results indicate that individuals with negative ΔCPM values exhibited smaller PAG grey matter volumes and larger FMC grey matter volumes. Exploratory analyses with PAG grey matter volume estimates revealed a negative correlation between PAG grey matter volume and LF-HRV (r= −0.54; p= 0.023).

**Figure 4.**
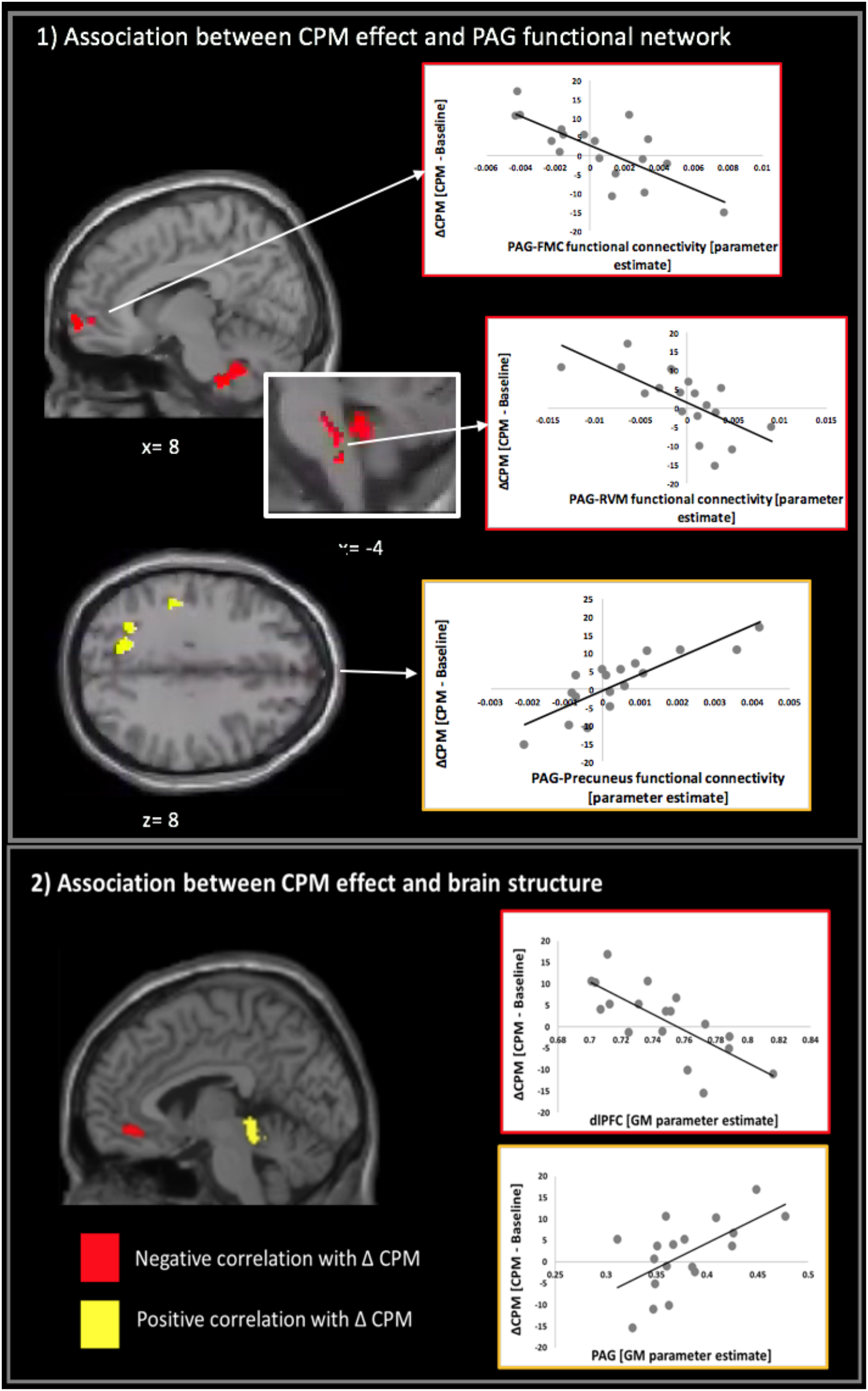
Association between CPM and brain structure and function. (1) Functional connectivity between the PAG and both the FMC and RVM was negatively associated with ΔCPM Cold-ON [CPM Cold-ON– Baseline pressure]. (2) ΔCPM Cold-ON was negatively associated with FMC grey matter volume and positively associated with PAG grey matter volume.

We did not observe any association between grey matter volume and LF-HRV during cold, nor between grey matter volume and the ΔCPM x LF-HRV interaction term.

#### Exploratory analyses of the association between PAG FC and pressure pain threshold and cold-pain ratings

We did not observe any positive or negative association between gray matter volume and baseline pressure pain ratings.

**TABLE 1.** Associations between CPM scores, autonomic reactivity during pain, functional connectivity and brain structure.

| Contra | Brain area | Direction of correlation | Cluster |  | Voxel |  |
| --- | --- | --- | --- | --- | --- | --- |
|  |  |  | *k* | *p FWE* | *T* | *MNI xyz* |
| Rs-fMRI regression analyses |  |  |  |  |  |  |
| *Association between FC and ΔCPM (Cold ON)* |  |  |  |  |  |  |
|  | Precuneus, left | positive | 195 | 0.034 | 6.12 | -18 -64 26 |
|  | Anterior supramarginal gyrus, right | positive | 225 | 0.019 | 5.26 | -44 -32 38 |
|  | Frontal lobe/FMC | negative | 672 | 0.008 * | 3.25 | -14 52 6 |
|  | Rostral ventro-medial medulla | negative | 489 | 0.039 * | 4.09 | 6 -36 -48 |
| *Association between FC and LF-HRV during cold pain* |  |  |  |  |  |  |
|  | Angular gyrus, right | negative | 930 | 0.001* | 5.24 | 66 -16 -24 |
| VBM regression analyses |  |  |  |  |  |  |
| *Association between GM volume and ΔCPM (Cold ON)* |  |  |  |  |  |  |
|  | FMC | negative | 85 | 0.027# | 4.57 | 2 39 -12 |
|  | PAG | positive | 10 | 0.045#* | 3.26 | -2 -33 -8 |

**FMC = Frontal medial cortex; PAG= periaqueductal grey; \* results obtained with a cluster forming threshold $p < 0.005$ ; # results obtained with a small volume correction.**

### Correlation between CPM associated PAG functional connectivity estimates, grey matter volume and LF-HRV

We performed post-hoc correlation analyses to explore associations between CPM related brain areas and LF-HRV. We observed a positive correlation between the strength of both PAG-FMC and PAG-RVM FC with LF-HRV, indicating that individuals with the highest increase in LF-HRV during pain had also the strongest FC among these the PAG, RVM and FMC (Figure 5, A and B). We also observed a negative correlation between PAG grey matter volume and LF-HRV during cold pain, indicating that individuals with the smallest PAG grey matter volumes had the highest increases in LF-HRV during pain (Figure 5, C).

**Figure 5.**
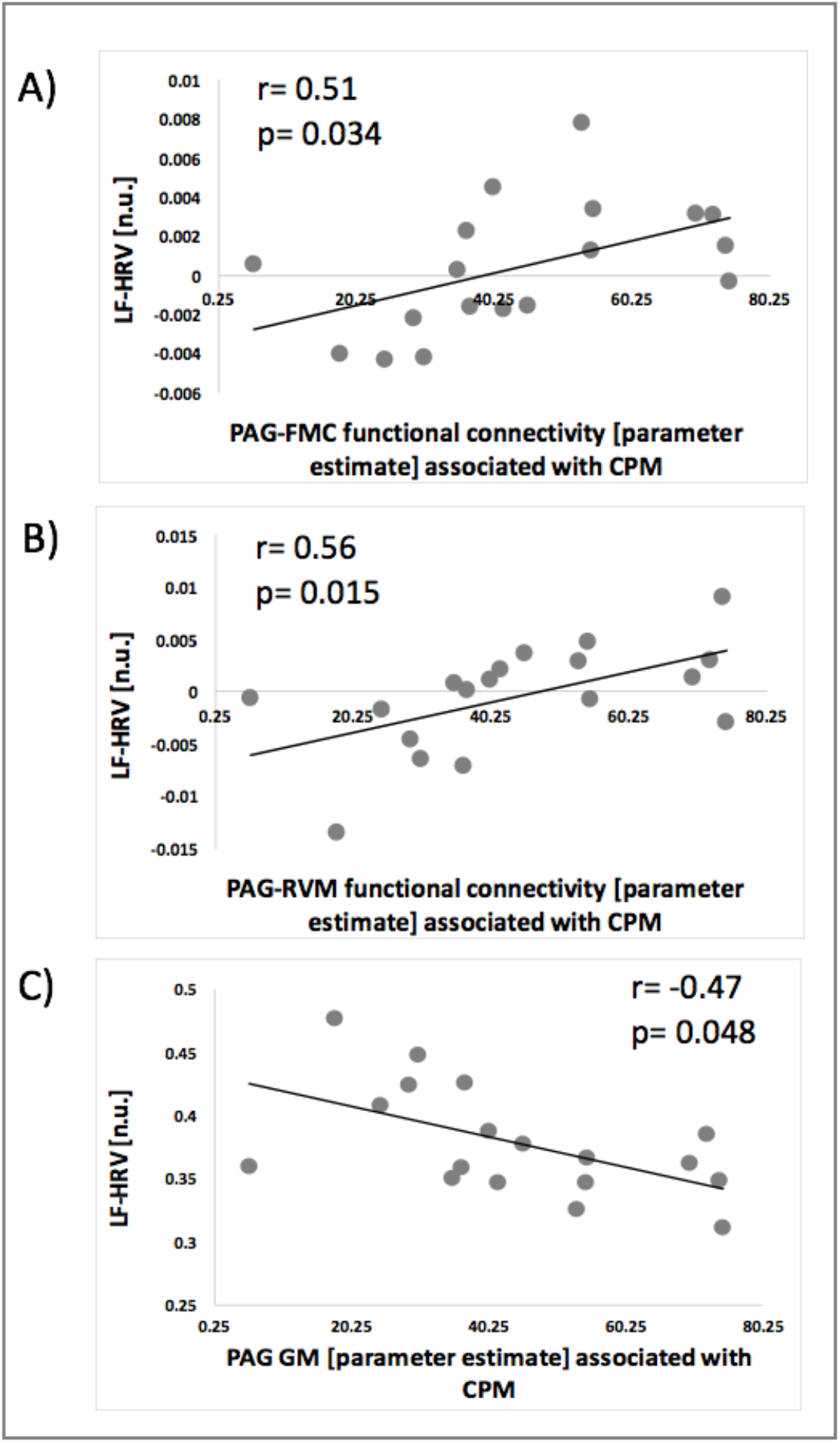
Correlation between LF-HRV and parameter estimates of CPM associated FC between PAG and FMC (A), PAG and RVM (B) and PAG grey matter volume.

### Mediation analysis

We observed a mediating role of PAG-FMC FC in the relationship between LF-HRV and the CPM effect. The total effect of LF-HRV on ΔCPM was significant (path C; t = −2.23, p = .041). Adding PAG-FMC FC as the mediator, the direct effect of LF-HRV on ΔCPM was no longer significant (path C’; t <1, p = .31), whereas the indirect path via PAG-FMC FC was significant (95% CI [−0.27, −0.08]). Therefore, the effect of LF-HRV on ΔCPM was largely mediated by PAG-FMC FC (Figure 6).

**Figure 6.**
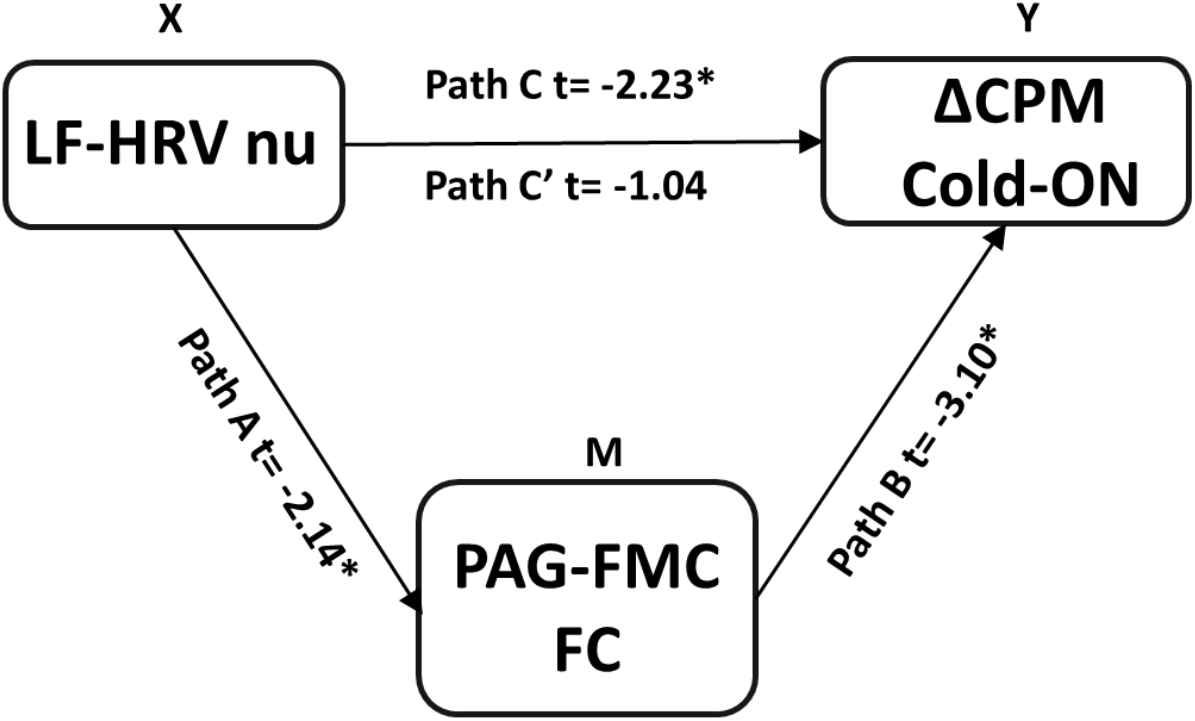
Mediation analysis indicated that the association between autonomic reactivity during cold pain (LF-HRV) and the efficiency of the descending inhibitory pain mechanisms (ΔCPM) was strongly mediated by left PFC-FMC. The t value for each path is presented. *p<.05.

## DISCUSSION

ANS activity can influence pain perception (Saccò et al., 2013). This interaction might involve descending pain modulatory pathways. In this healthy volunteer study, we used restingstate fMRI and voxel-based morphometry to investigate brain areas underlying one’s ability to engage descending pain modulatory mechanisms (as measured by CPM) and autonomic reactivity to cold pain (as measured by the LF-HRV index). We observed an association between LF-HRV during pain and the magnitude of an individual’s CPM response. The CPM response was associated with PAG-FMC and PAG-RVM FC, and with PAG and FMC grey matter volume. The same brain areas were also associated with LF-HRV during pain. The strength of the association between CPM and LF-HRV was mediated by PAG-FMC FC in response to tonic cold pain. Our results provide evidence for the mechanisms through which central, peripheral and pain systems interact, indicating that an adaptive autonomic reaction to pain is involved in descending pain inhibition.

We have demonstrated an association between each person’s autonomic reactivity to pain and the efficiency of their descending pain control systems, namely a negative association between an LF-HRV during tonic pain and a decrease in subjective ratings of pressure pain during simultaneous application of a cold pain. HRV is a measure of the interplay between sympathetic and parasympathetic activity in autonomic control of the heart. Whereas HF-HRV is purely a measure of the parasympathetic influence, LF-HRV (especially the normalized measure) reflects an influence of both the sympathetic and parasympathetic nervous systems (Reyes del Paso et al., 2013), accordingly providing a more generalized index of autonomic balance (Burr, 2007; Eickhoff, Bzdok, Laird, Kurth, & Fox, 2012; Goldstein et al., 2011; Moak et al., 2007; Rahman et al., 2011). LF-HRV has been associated with lower ratings of unpleasantness to heat pain stimulation and higher thresholds for barely noticeable and moderate pain (Appelhans & Luecken, 2008). Here, we went one step further by investigating the relationship of ANS reactivity to pain, with a focus on descending pain control, as indexed by CPM scores. Our findings accord with one previous study in patients with fibromyalgia showing that systolic blood pressure responses during the cold pressor test are positively associated with the CPM inhibiting effect, suggesting that chronic pain might be linked to a reduced blood pressure response to the conditioning stimulus during CPM (Chayale et al., 2014). Our study, in healthy participants, supports the existence of a functional pain inhibitory feedback system described in animal models that is responsible for blood pressure-related hypoalgesia (Ghione, 1996). This model describes a reflexive increase in sympathetic arousal during pain, resulting in an increase in blood pressure (Reis, Ruggiero, & Morrison, 1989). The increase in blood pressure stimulates baroreceptors, whose afferent information activates descending pain modulatory pathways, modulating perceived pain and regaining homeostasis (Bruehl & Chung, 2004). In this work, we focused our interest on the relationship between ANS reactivity to a tonic cold painful stimulus and CPM response. However, both test and conditioning stimuli are likely to produce acute ANS reactions. Future studies should consider not only ANS reactions to conditioning stimuli, and test stimuli in isolation, but also potential effects of combined ANS reactions elicited during simultaneous presentations of these stimuli and their relationship to the CPM response.

Our next aim was to investigate relationships between CPM response, brain FC and brain structure, and to investigate whether the HRV-CPM relationship is mediated by a common neural network. For the FC analyses, we considered the PAG as our seed region. The PAG is considered a core hub for serotoninergic neurons involved in descending pain modulation that project to the RVM, the output pathway for pain control to the dorsal horn of the spinal cord (Fields, 1999; Mtui, Gruener, & FitzGerald, 2011). The PAG also receives cortical projections, originating principally from the FMC/PFC (An, Bandler, Öngür, & Price, 1998; Neafsey, Hurley-Gius, & Arvanitis, 1986); stimulation of these cortical areas can induce analgesia (Benarroch, 2012). We observed that participants who reported a reduction in pressure pain in the presence of a tonic painful conditioning stimulus, also had increased FC strength between the PAG and both the FMC and RVM. Other fMRI studies in humans investigating brain activity underpinning CPM have provided contrasting results. In a study by Piché and colleagues (Piché, Arsenault, & Rainville, 2009), a sustained increase in orbito-frontal cortex activity during the application of a cold contralateral conditioning stimulus was associated with a decrease in subjective response to evoked painful electrocutaneous test stimuli. Similarly, Bogdanov and colleagues described that individual differences in CPM (i.e. reduction in painful laser heat ratings) related to levels of PFC activation in response to the early part of the cold conditioning stimulus (Bogdanov et al., 2015). Youssef and colleagues, on the other hand, suggested that engagement of the prefrontal and cingulate areas interferes with the generation of CPM (Youssef et al., 2016). In contrast to the studies of Piche and Bogdanov, the conditioning stimulus adopted in this later study was hypertonic saline-induced muscle pain, suggesting that the discordance in results between studies is likely to relate to the specific pain modalities and/or stimulation methodology adopted in the CPM investigation (Kemp et al., 2019; Kennedy, Kemp, Ridout, Yarnitsky, & Rice, 2016). In a resting-state fMRI study by Harper and colleagues, higher FC between the PAG, left insula and perigenual ACC was associated with more efficient inhibitory CPM responses in both healthy participants and patients with fibromyalgia, whereas PAG connectivity with the dorsal pons was associated with greater CPM inhibition only in the control group. Greater PAG connectivity to the caudal pons/rostral medulla was associated with descending pain inhibition in healthy participants, but with pain facilitation in fibromyalgia patients (Harper et al., 2018). Here, we provide new knowledge to the field by showing that the same brain structures are associated with both the cardiovascular responses to the conditioning stimulus and the efficiency of descending pain control. Our data indicate that the strength of FC between these structures is a mediator of the ANS-CPM association. Dysregulation of the three-way interaction between brain FC, efficiency of descending pain control mechanisms and autonomic reactivity might explain various symptoms of chronic pain conditions such as low HRV and impaired descending pain control compared to healthy controls (Lewis, Rice, & McNair, 2012; Tracy et al., 2016). Future studies are required to investigate the existence and dysregulation of such a mechanism in chronic pain conditions.

Brain structure was also associated with the CPM effect. Specifically, individuals with larger FMC and smaller PAG volumes exhibited an inhibitory CPM effect. Previous reports have described associations between grey matter volume reductions in the orbito-frontal cortex and the FMC (together with other areas such as the thalamus, insula, posterior cingulate cortex) and increased sensitivity to rectal pain stimuli in healthy volunteer participants (Elsenbruch et al., 2014). Reduction in FMC grey matter is found in patients with chronic complex regional pain syndrome (Geha et al., 2008), fibromyalgia (Kuchinad et al., 2007) and chronic back pain (Fritz et al., 2016). Similarly, patients with functional dyspepsia have decreased cortical thickness in prefrontal and dorso-lateral areas (Liu et al., 2018). A negative correlation was described between pain and dlPFC volumes in patients with chronic complex regional pain syndrome (Barad, Ueno, Younger, Chatterjee, & Mackey, 2014), whereas pain thresholds were correlated positively with dlPFC volumes in patients with chronic myofascial pain (Niddam, Lee, Su, & Chan, 2017). Here, our results show that individuals with larger FMC volumes report more efficient descending pain inhibition, whereas individuals with smaller grey matter volumes in this area showed pain facilitation during the application of the conditioning stimulus. Taken together these findings engender the working hypothesis that PFC volume reductions in multiple chronic pain conditions may be associated with impairments in descending pain control. Our data suggest that individuals with smaller FMC volumes exhibited pain facilitation (i.e. an increase in pain with the contralateral presentation of the conditioning stimulus), whereas those with larger FMC volumes exhibited a reduction in pain (indicating efficient descending pain modulating mechanisms). We speculate that, in chronic pain patients, a reduction in FMC/PFC volume might be involved in the phenotypic switch between descending inhibition to pathological pain facilitation (Potvin & Marchand, 2016).

PAG volumes correlated positively with CPM response. Individuals with larger PAG volumes reported pain facilitation, whereas comparatively smaller PAG grey matter volumes were associated with pain inhibition. PAG grey matter volumes were also associated with the HRV responses to cold pain, where individuals with smaller PAG volumes reported the strongest increase in HRV in response to pain. Reports of PAG grey matter alterations and their involvement in pain chronification are contrasting. Reductions in PAG grey matter volumes have been described in fibromyalgia patients and have been further related to pain facilitation effects during a CPM paradigm (Harper et al., 2018). Other studies, however, reported an increase in PAG grey matter volume in patients suffering from medication-overuse headache (Riederer et al., 2013) and migraine (Rocca et al., 2006). A decrease in PAG grey matter volume after a significant improvement of clinical symptoms in medication-overuse headache patients was also observed (Riederer et al., 2013). The reasons for these contrasting reports are currently unclear. They might reflect methodological difficulties in accurately measuring volumes of small structures such as the PAG, by using MRI methodology which usually relies on cortically biased methods. Presumably, both decreases and increases in PAG grey matter volume might be dysfunctional under specific circumstances; PAG grey matter volume changes might be related to inefficient pain inhibition, perhaps the consequences of maladaptive neuronal plasticity increasingly accepted to occur in chronic pain states (May, 2008). Here, we suggest that the pain facilitatory/inhibitory effect of the PAG might be related to the degree of ANS input. Speculatively, this mechanism might involve the baroreceptors, signaling the state of body arousal and blood pressure via the vagus and glossopharyngeal nerve, to the solitary nucleus of the medulla and the PAG. In line with this theory, we hypothesize that conditions such as hypertension-related hypoalgesia (increased pain thresholds in individuals with high blood pressure) might involve over-engagement of descending pain inhibitory mechanisms. The opposite may also be true; individuals with chronic low blood pressure are prone to thermal hyperalgesia (Duschek, Dietel, Schandry, & del Paso, 2009; Duschek, Schwarzkopf, & Schandry, 2008), which we speculate may involve a shift toward descending pain facilitation.

In conclusion, this study combined rs-fMRI, recording of the ANS responses to pain and assessments of descending pain control, providing mechanistic insights into the association between the ANS and pain in the brain. Our data adds to the large body of research focusing on the link between the individual differences in autonomic balance and physical and emotional well-being, underpinned by the PFC (Thayer et al., 2009). Associations identified between CPM response, resting-state connectivity networks involved in descending pain control and flexibility in heart rate variability suggest that individuals with stronger engagement of the ANS in response to pain are also more efficient in triggering descending pain inhibition, which may lead to more expedient recovery of homeostasis. Future studies will shed light on potential dysregulation of these mechanisms in patients with chronic painful conditions, which in turn may signpost future therapeutic interventions for these patients. Therapies focused on stimulation of the ANS (such as HRV biofeedback, or baroreceptor stimulation) might help to re-gain appropriate levels of control over not only the ANS, but also mechanisms of endogenous pain control.

## Acknowledgments

This work was funded by a Medical Research Council Experimental Medicine Challenge Grant (MR/N026969/1) and by the EFIC-Grünenthal-Grant (EGG) 2018 awarded to EM. MAH, SM and SW are also supported by the NIHR Biomedical Research Centre for Mental Health at the South London and Maudsley NHS Trust. We also wish to thank the Wellcome Trust for their ongoing support of our neuroimaging research and Simon Hill for both the development and calibration of the stimulation equipment and Giovanni Calcagnini for his input on Heart Rate Variability analysis.

